# Allele-specific silencing of a dominant *SETX* mutation in familial amyotrophic lateral sclerosis type 4

**DOI:** 10.1101/2024.10.11.617871

**Authors:** Audrey Winkelsas, Athena Apfel, Brian Johnson, George Harmison, Dongjun Li, Vivian Cheung, Christopher Grunseich

## Abstract

Amyotrophic lateral sclerosis 4 (ALS4) is an autosomal dominant motor neuron disease that is molecularly characterized by reduced R-loop levels and caused by pathogenic variants in *senataxin* (*SETX*). *SETX* encodes an RNA/DNA helicase that resolves three-stranded nucleic acid structures called R-loops. Currently, there are no disease-modifying therapies available for ALS4. Given that *SETX* is haplosufficient, removing the product of the mutated allele presents a potential therapeutic strategy. We designed a series of siRNAs to selectively target the RNA transcript from the ALS4 allele containing the c.1166T>C mutation (p.Leu389Ser). Transfection of HEK293 cells with siRNA and plasmids encoding either wild-type or mutant (Leu389Ser) epitope tagged SETX revealed that three siRNAs specifically reduced mutant SETX protein levels without affecting the wild-type SETX protein. In ALS4 primary fibroblasts, siRNA treatment silenced the endogenous mutant *SETX* allele, while sparing the wild-type allele, and restored R-loop levels in patient cells. Our findings demonstrate that mutant *SETX*, differing from wild-type by a single nucleotide, can be effectively and specifically silenced by RNA interference, highlighting the potential of allele-specific siRNA as a therapeutic approach for ALS4.

## INTRODUCTION

Amyotrophic lateral sclerosis 4 (ALS4) is an autosomal dominant motor neuron disease with an average age of onset in the teenage years. Most patients have a slowly progressive disease course that includes symmetric muscle weakness in a predominantly distal distribution with hyperreflexia.^1^ There is currently no disease modifying therapy available.

ALS4 is caused by heterozygous missense mutations in *senataxin* (*SETX*).^2^ Senataxin, an ortholog of the yeast sen1, is an ATP-dependent RNA/DNA helicase that resolves R-loops, three-stranded nucleic acid structures that include RNA/DNA hybrids and the displaced single-stranded DNA.^3–5^ We have previously shown that autosomal dominant mutations in senataxin lead to gain of the SETX helicase function^6^ and a reduction in R-loops.

One of the common ALS4 mutations is the c.1166T>C transition (p.Leu389Ser, L389S). A Maryland family with an eleven-generation history of juvenile ALS harbors this *SETX* mutation, and many individuals from this family are followed by our group.^7,8^ Previously, we performed deep phenotyping of these patients and biomarker identification, as well as extensive studies of the molecular effects of ALS4 mutations on SETX function.^6,8^

Silencing the mutated allele may be a viable therapeutic strategy for ALS4 since the *SETX* gene is haplosufficient. Small interfering RNAs (siRNAs) are tools for sequence-based gene silencing and have been shown to offer allele specificity. Here, we identified and characterized siRNAs against the L389S *SETX* allele and demonstrated their effects on SETX biology and the functioning of patient cells.

## RESULTS

### siRNAs that target the L389S *SETX* mutation that causes ALS4

We designed a series of siRNAs to target the L389S-encoding allele in which the nucleotide complementary to the mutation is located at a different position within the siRNA sequence (Figure S1). The candidate siRNAs were scored based on preferred sequence characteristics, such as GC content and the base composition at specific positions within the siRNA.^9^ We selected three of the top-scoring siRNAs (siRNAs 1.02, 1.11, and 1.16) to test experimentally.

Additional nucleotide mismatches into siRNAs can enhance allelic discrimination,^10^ so we also included an siRNA with a second mismatch to the wild-type allele. The double mismatch siRNA, named 2.11, was designed using *in silico* siRNA efficacy prediction tool.^11^

### Assessing the siRNAs using a reporter system

To evaluate the efficacy of the siRNAs, we transfected HEK293 cells with siRNAs and a plasmid encoding either wild-type SETX or L389S SETX, each fused to an N-terminal Halo tag. We then quantified the Halo-SETX fusion protein in the cell lysates with immunoblotting. All three siRNAs (1.02, 1.11, 1.16) significantly reduced the levels of mutant SETX (Bonferroni corrected p < 0.0001, one-way ANOVA) to 28%, 27%, and 30%, respectively (Figures 1A and 1B). Likewise, the double-mismatch siRNA 2.11 reduced mutant SETX levels to 26% (Bonferroni corrected p = 0.0038, one-way ANOVA, Figures 1C and 1D). Notably, siRNAs 1.11, 1.16, and 2.11 specifically targeted the mutant SETX without affecting wild-type SETX expression, while siRNA 1.02 decreased wild-type SETX expression to 38% (Bonferroni corrected p < 0.001, one-way ANOVA).

**Figure 1.**
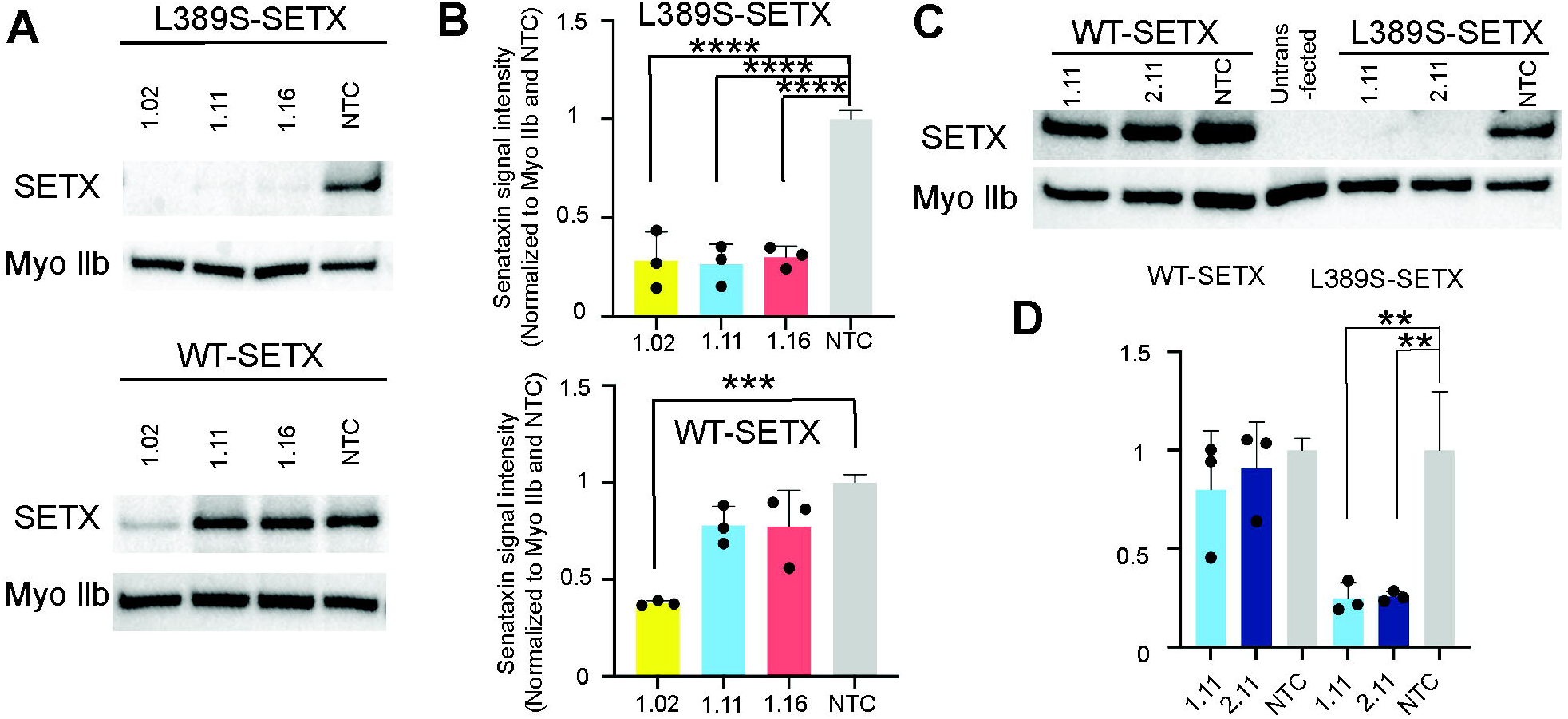
Effective and preferential knockdown of the L389S-SETX with siRNA. (A) HEK293 cells were transfected with three different *SETX*-targeting siRNA (1.02, 1.11, 1.16) and plasmid encoding either mutant (L389S) or wild-type (WT) Halo-SETX fusion proteins. SETX protein levels were assessed by Western blotting with Myo IIb used as a loading control. (B) Quantification of SETX protein knockdown showing allele specific efficacy for siRNA 1.11 and 1.16, and non-specific efficacy for 1.02. (C+D) HEK293 cells transfected with siRNA containing an additional nucleotide mismatch along with the Halo-SETX fusion proteins shows evidence of allele specificity with significant knockdown of the mutant, but not wild-type, allele. ** p < 0.01, *** p < 0.001, **** p < 0.0001.

### Allele-specific and dose-dependent knockdown of endogenous, mutant *SETX*

Next, we assessed whether the siRNAs affect the expression of *SETX* in cells from patients with ALS4. We transfected siRNAs 1.11, 1.16, and 2.11 into the cultured primary fibroblasts and measured the expression of the endogenous mutant and wild-type alleles using quantitative RT-PCR with probes that discriminate between cDNA from the two alleles (Figure S2A). There was a dose-dependent decrease in the amount of mutant *SETX* (Figure S2B). All three siRNAs achieved similar potency and specificity (Figure 2A). In cells from three individuals with ALS4, 25nM of each siRNA (siRNA 1.16, 1.11, and 2.11) knocked down the mutant allele from an average of 52% of total transcript abundance to 11% (p = 0.003), 15% (p = 0.009), and 12% (p = 0.003), respectively (Figure S2B). In control fibroblasts without the c.1166T>C mutation, siRNA 2.11 showed no significant reduction in wild-type *SETX* mRNA levels, further confirming its specificity for the mutant *SETX*. Sanger cDNA sequencing from two additional patient fibroblast lines also demonstrated that siRNAs 1.11, 1.16, and 2.11 knocked down the mutant allele (C-bearing) relative to the wildtype (T-bearing) allele (Figures 2B and S3A).

**Figure 2.**
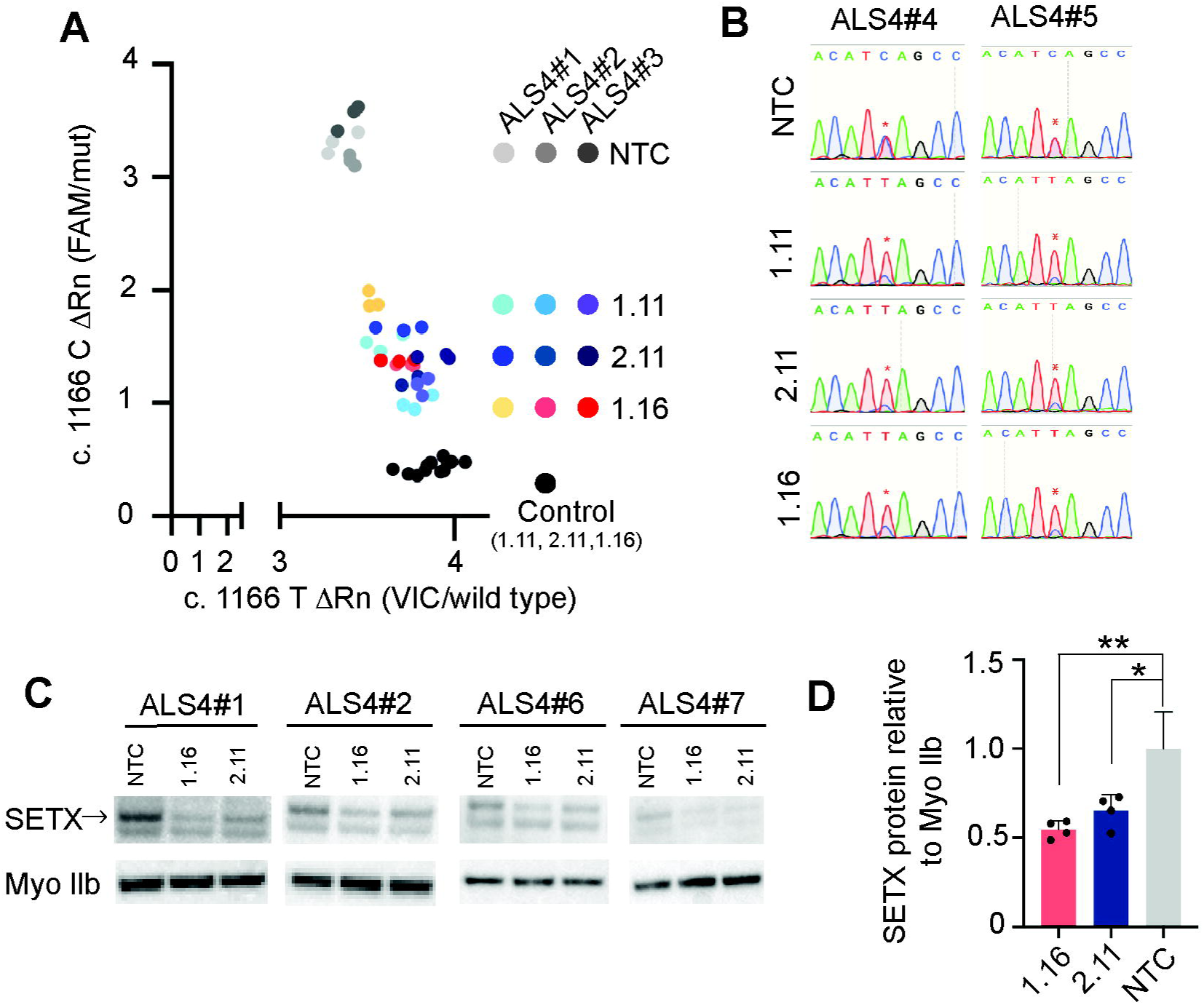
siRNA efficacy in ALS4 derived patient fibroblasts. (A) Mutant *SETX* transcript is knocked down in ALS4 fibroblasts with siRNAs 1.11, 2.11, 1.16, and non-targeting control (NTC). RNA was collected at 72 hrs post-transfection and transcript levels from the wild-type (x-axis) and mutant (y-axis) alleles were measured by quantitative RT-PCR and *SETX* SNP genotyping assay. Control is a healthy control fibroblast. (B) Sanger sequencing of cDNA from ALS4 treated fibroblasts showing the relative abundance of the mutant allele (C-bearing) to wild-type allele (T-bearing). (C+D) Western blot analysis of lysate from fibroblasts treated with siRNA for 3 days, probed with antibodies for SETX and Myo IIb (loading control), showing a significant reduction in SETX protein in the siRNA 1.16 and 2.11 treated samples. * p < 0.05, ** p < 0.01.

We further evaluated the impact of the siRNAs on SETX protein expression by transfecting fibroblasts from four patients. On average, SETX protein levels were reduced by 35% (p = 0.01) with siRNA 2.11 and 45% (p = 0.0019) with siRNA 1.16 (one-way ANOVA, Figures 2C and 2D). These findings indicate that the siRNAs are effectively reducing both senataxin transcript and protein expression.

### Effects of siRNAs on SETX gain of function activity

Previously, we showed that the L389S mutation in senataxin leads to fewer R-loops. Here, we asked if silencing the *SETX* mutant allele increases R-loop abundance. Primary fibroblasts from patients and controls were transfected with siRNA 2.11. After confirming allele-specific knockdown, we carried out R-loop enrichment with S9.6 antibody in DNA-RNA immunoprecipitation.^12^ We then measured R-loop abundance at four known R-loop regions, including the well-characterized *ACTB* and *BAMBI*. For all four R-loop regions tested, R-loop abundance increased significantly in patient cells treated with siRNA 2.11 (*ACTB, BAMBI, RPL13A*: p < 0.0001, *AANCR*: p = 0.0019). In our previous study, we found that the reduction in R-loop abundance in *BAMBI* promoter led to more methylation and silencing of *BAMBI*, a negator of TGFβ, and thus activation of TGFβ. Here, we found that silencing of the mutant *SETX* allele led to a greater than 50% increase in R-loop at the *BAMBI* promoter. Together, the results show that allele-specific silencing of the *SETX* mutation restores the R-loop defects in cells from ALS4 patients.

## DISCUSSION

ALS4, an autosomal dominant motor neuron disease caused by heterozygous mutations in senataxin, presents many of the challenges common to rare diseases in therapeutic development. In this study, we applied RNA interference (RNAi) to selectively silence the mutant senataxin allele while maintaining expression of the wild-type allele. We identified three siRNAs (1.11, 1.16, and 2.11) that specifically target the L389S mutation in SETX, which is responsible for ALS4. Silencing the mutant allele restored normal senataxin helicase activity.

Previously, we demonstrated that autosomal dominant mutations in senataxin result in a gain of helicase function, leading to a reduction in R-loop levels in patient cells. This includes decreased R-loop abundance at the promoter of *BAMBI*, a negative regulator of TGFβ.^6,8,13^ Here, we show that allele-specific silencing of the mutant *SETX* copy increases R-loop levels, including those at the *BAMBI* promoter, providing further evidence of restored RNA/DNA helicase function in patient-derived cells.

Targeted knockdown of the mutant senataxin allele is crucial, as silencing both *SETX* alleles would likely replicate the loss-of-function phenotype seen in patients with oculomotor apraxia type 2 (AOA2), an autosomal recessive disorder caused by homozygous *SETX* mutations. Carriers of the autosomal recessive disorder AOA2, who possess heterozygous loss-of-function *SETX* mutations, are unaffected. This suggests that selective knockdown of the mutant allele, while maintaining wild-type *SETX* expression, may offer therapeutic benefit in ALS4.

An advantage of siRNA technology is its ability to design therapeutic agents that specifically target the gene of interest. siRNA therapies were first approved by the US Food and Drug Administration (FDA) in 2018 for the treatment of transthyretin-associated amyloidosis,^14^ and since then, several siRNAs have received FDA approval, with many more currently in clinical trials.^15^ By targeting the underlying genetic mechanisms, siRNA therapies offer a precise way to intervene in the disease process for various Mendelian disorders.^16^ The ability to design gene-targeting therapies with specificity has made this approach feasible for treating rare neurological diseases.^17^ Our findings demonstrate that this gene-targeting strategy can mitigate toxicity in a rare form of genetic ALS. Importantly, our approach can be adapted for allele-specificity, enabling selective knockdown of the mutant *SETX* allele with minimal impact on the wild-type copy.

In summary, this study underscores the potential of allele-specific siRNA as a therapeutic approach for ALS4. Our experiments, conducted in seven patient samples, provide robust evidence of efficacy across multiple ALS4 patients, supporting the further development of this strategy for clinical use. The next step involves optimizing the siRNAs, such as through chemical modifications, to enhance pharmacodynamic properties like half-life, uptake, and activity, thus improving therapeutic efficacy^18^ and advancing this ALS4 therapy toward clinical trials.

## MATERIALS AND METHODS

### Patients and cell culture

ALS4 patients were identified and confirmed by genetic testing to have the *SETX* mutation at codon L389S. ALS4 patients in this study included males aged 27, 31, 33, 36, and 46 years old and females aged 52 and 61 years old. All skin biopsies were performed at the National Institutes of Health (NIH) in Bethesda, MD under IRB-approved protocol 00-N-0043 “Clinical and Molecular Manifestations of Inherited Neurological Disorders” and 20-N-0064 “An observational study to assess clinical manifestations and biomarkers in amyotrophic lateral sclerosis type 4 and other inherited neurological disorders of RNA processing.” Written informed consent was received from all participants before study inclusion. Dermal skin fibroblasts were obtained by 3 mm punch biopsy of the anterior forearm and subsequently expanded in media containing DMEM with 10% fetal bovine serum and passaged at a 1:3 splitting ratio using Trypsin-EDTA (0.25%) and grown at 37 °C with 5% CO2. HEK293 cells were expanded in media containing DMEM with 10% fetal bovine serum.

### siRNA

siRNAs were obtained from Dharmacon (standard processing, desalted and deprotected duplex) and designed to achieve allele specificity against the mutant allele of *SETX*. siRNAs in this study contained the following sense sequences, which include UU overhangs at the 3’ end: 1.02- 5’ UGAAGAAAUGGAAACAUCAUU 3’, 1.11- 5’ GGAAACAUCAGCCAGUGUAUU 3’, 1.16- 5’ CAUCAGCCAGUGUACUUCAUU 3’, 2.11- 5’ GGAAACAUCAGCUAGUGUAUU 3’. The non-targeting control (NTC) siRNA was designed against the target sequence 5’ UAAGGCUAUGAAGAGAUAC 3’. HEK293 cells were seeded 24 hours before transfection in DMEM, 10% FBS without antibiotics. Cells were transfected with 25 nM siRNA using RNAiMAX reagent (Invitrogen) and subsequently transfected with Halo-SETX fusion constructs using lipofectamine 3000 (Invitrogen). Media was changed 24 hours after transfection and cells were harvested 72 hours post transfection for Western blot analysis. Human fibroblasts were seeded 24 hours before transfection in media without antibiotics and subsequently transfected with 25 nM siRNA in RNAiMAX reagent. Media was changed the following day and cells were harvested after 72 hours.

### Western blot analysis

Fibroblasts and HEK293 cells were lysed with RIPA buffer (50 mM Tris, pH 8, 150 mM NaCl, 1% NP-40, 0.5% sodium deoxycholate, 0.1% SDS). Western blots were performed with antibodies for SETX (Bethyl) and myosin IIb (Cell Signaling Technology).

### Messenger RNA expression analysis

Cells were homogenized in 500 mL TRIzol reagent (Invitrogen) and centrifuged for 10 min at 4 °C. 100 mL chloroform was added to the supernatant, and the mixture was vortexed and spun down again. The aqueous phase was transferred to a separate tube and purified using the RNA-clean up kit (Qiagen). 1 ug total RNA was converted to cDNA using the High Capacity cDNA Reverse Transcriptase kit (Invitrogen). Transcript levels from the wild-type and mutant alleles were measured using quantitative RT-PCR analysis with a *SETX* c.1166 T>C SNP genotyping probe (Invitrogen). Each reaction contained 2.5 uL of TaqMan genotyping master mix, 1.375uL of water, 0.125 uL of 40X *SETX* genotyping probe (Invitrogen) and 1 uL of cDNA. Standard curves were generated by mixing the ALS4 and control cDNA using the following ratios of input ALS4:control cDNA; 1:9, 3:7, 1:1, 7:3, 9:1. Sanger sequencing was performed by PCR amplification of cDNA from treated human fibroblasts to detect the relative abundance of transcript from the mutant and wild-type alleles. Primers 5’ CTTTCTGGCCAGCGTTACACTG and 5’ GTGCTGTTATGAACACGCATG were used for PCR amplification and primer 5’ GTGATTCTGGATCGCCTTGG was used for sequencing.

### DNA-RNA hybrid immunoprecipitation (DRIP)

The immunoprecipitation procedure was adapted from previous studies. 5×10^6^ primary fibroblasts were lysed in 600 μL cell lysis buffer (50 mM PIPES, pH 8.0, 100 mM KCl, 0.5% NP-40), and nuclei were collected by centrifugation. The nuclei pellet was resuspended in 300 μL nuclear lysis buffer (25 mM Tris-HCl, pH 8.0, 1% SDS, 5 mM EDTA). Genomic DNA along with R-loops was then extracted with phenol:chloroform and precipitated using ethanol. Purified DNA was resuspended in IP dilution buffer (16.7 mM Tris-HCl, pH 8.0, 1 mM EDTA, 0.01% SDS, 1% Triton-X100, 167 mM NaCl) and sonicated for 5 minutes using a Bioruptor (high setting, cycles of 30sec on/30sec off) to fragments with an average size of 500 base pairs. 5 μg of S9.6 monoclonal antibody (gift from Dr. Stephen H. Leppla at NIH) or non-specific mouse IgG (Santa Cruz) was used for each immunoprecipitation. Input and precipitates were analyzed by quantitative PCR using the primers in the following table.

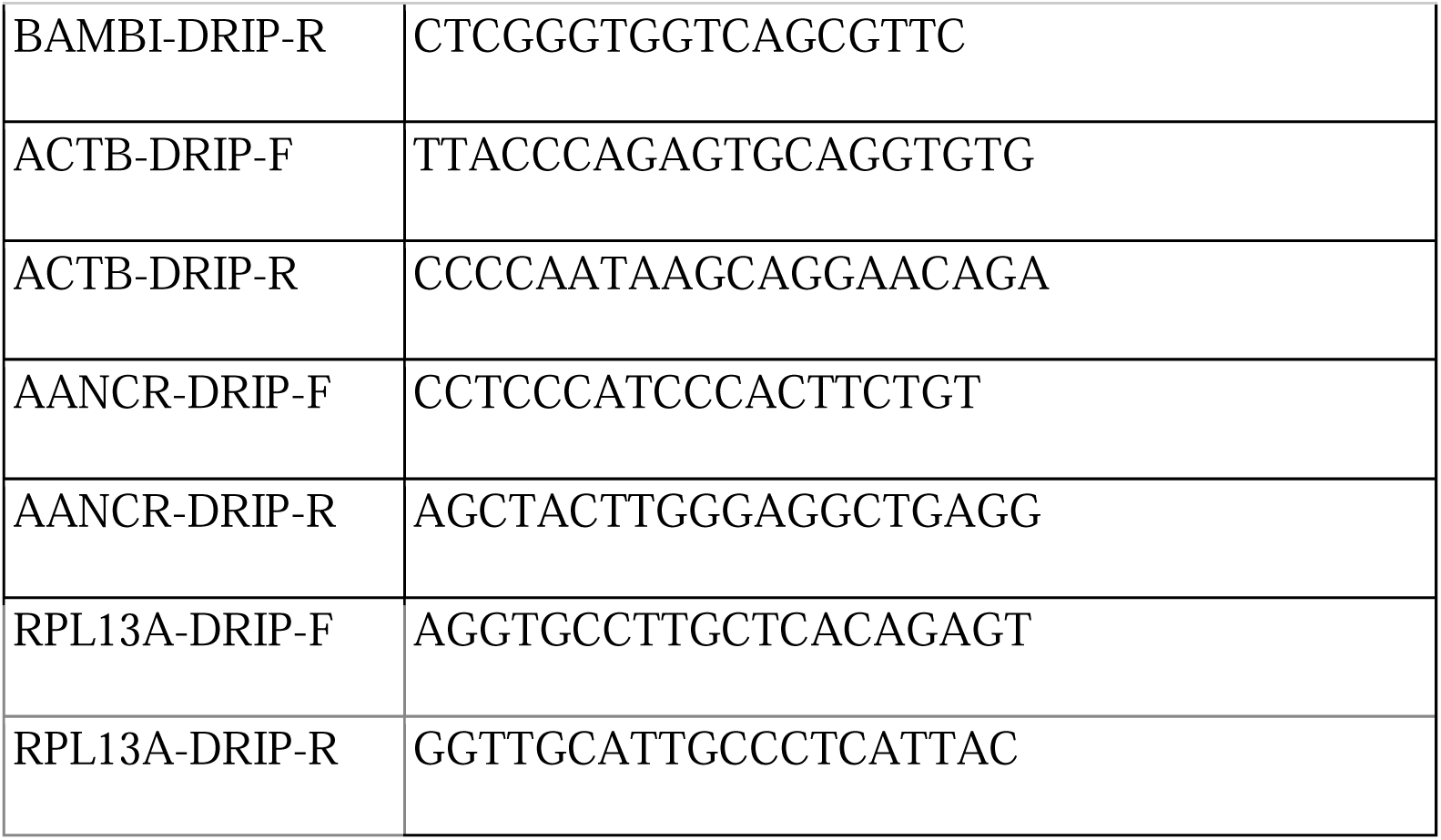

### Statistical Analysis

One-way ANOVA with Bonferroni’s multiple comparisons test was used to evaluate for statistical significance. Student t-test used for analysis of R-loop levels in figure 3. Error bars represent standard deviation.

**Figure 3.**
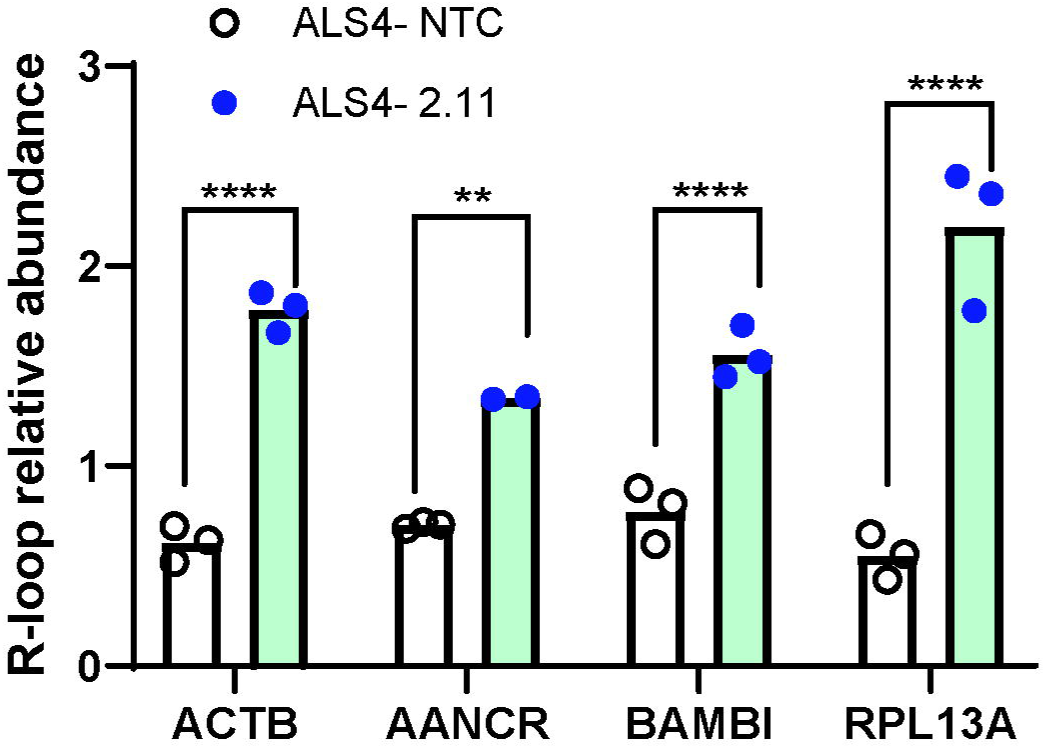
siRNA treatment corrects defects in patient R-loop levels. Primary fibroblasts from controls and patients were transfected with NTC or siRNA 2.11. After 72 hrs, DRIP with S9.6 was carried out, and R-loop abundance at *ACTB*, the enhancer RNA *AANCR*, *BAMBI* and *RPL13A* were measured by PCR. Results showed a significant increase in R-loop abundance at all four regions in the ALS4 fibroblasts. (** p < 0.01, **** p < 0.0001, t-test)

## DATA AVAILABILITY

All of the data in this study are available within the paper or can be obtained from the corresponding author upon request.

## ACKNOWLEDGEMENTS

This work is supported by intramural research funds from the National Institute of Neurological Disorders and Stroke, and extramural funds (ES034919 to VGC)

## AUTHOR CONTRIBUTIONS

Conceptualization: A.W., C.G., V.C.; Formal analysis: A.W., C.G., V.C.; Funding acquisition: C.G., V.C.; Investigation: all authors; Project management: A.W., C.G., V.C., Resources: C.G., V.C.; Supervision: C.G., V.C.; Writing-original draft: A.W., C.G., V.C.; Writing-review and editing: all authors.

---

**Supplemental figure 1. siRNA sequence diagram**

(A) Control and ALS4 sequence at coding sequence position 1166 highlighted in yellow with the mutant allele variant shown in red. siRNAs 1.02, 1.11, and 1.16 had high specificity scores and were chosen for further analysis. (B) siRNA 2.11 was generated with an additional base mismatch shown in blue to assist in allele specificity.

**Supplemental figure 2. siRNA characterization**

(A) Representative standard curve using the *SETX* c.1166 T>C SNP genotyping probe with a dilution series of the ALS4 and control fibroblast cDNA. (B) Dose-dependent knockdown of the mutant allele of L389S in an ALS4 patient fibroblast line in fibroblasts treated for 72 hrs with increasing doses of siRNA. (C) Quantification of knockdown of the mutant L389S *SETX* allele in three patient fibroblast lines (each indicated with a separate color circle) using 25 nM of siRNA.

**Supplemental figure 3.**

(A) Quantification of Sanger sequencing of cDNA from treated fibroblasts showing the relative abundance of the mutant allele (C-bearing) to the wild-type allele (T-bearing). Each circle represents an independent replicate.

