## Supplementary figures and images for "Allele-specific silencing of a dominant *SETX* mutation in familial amyotrophic lateral sclerosis type 4"

### Figure S1

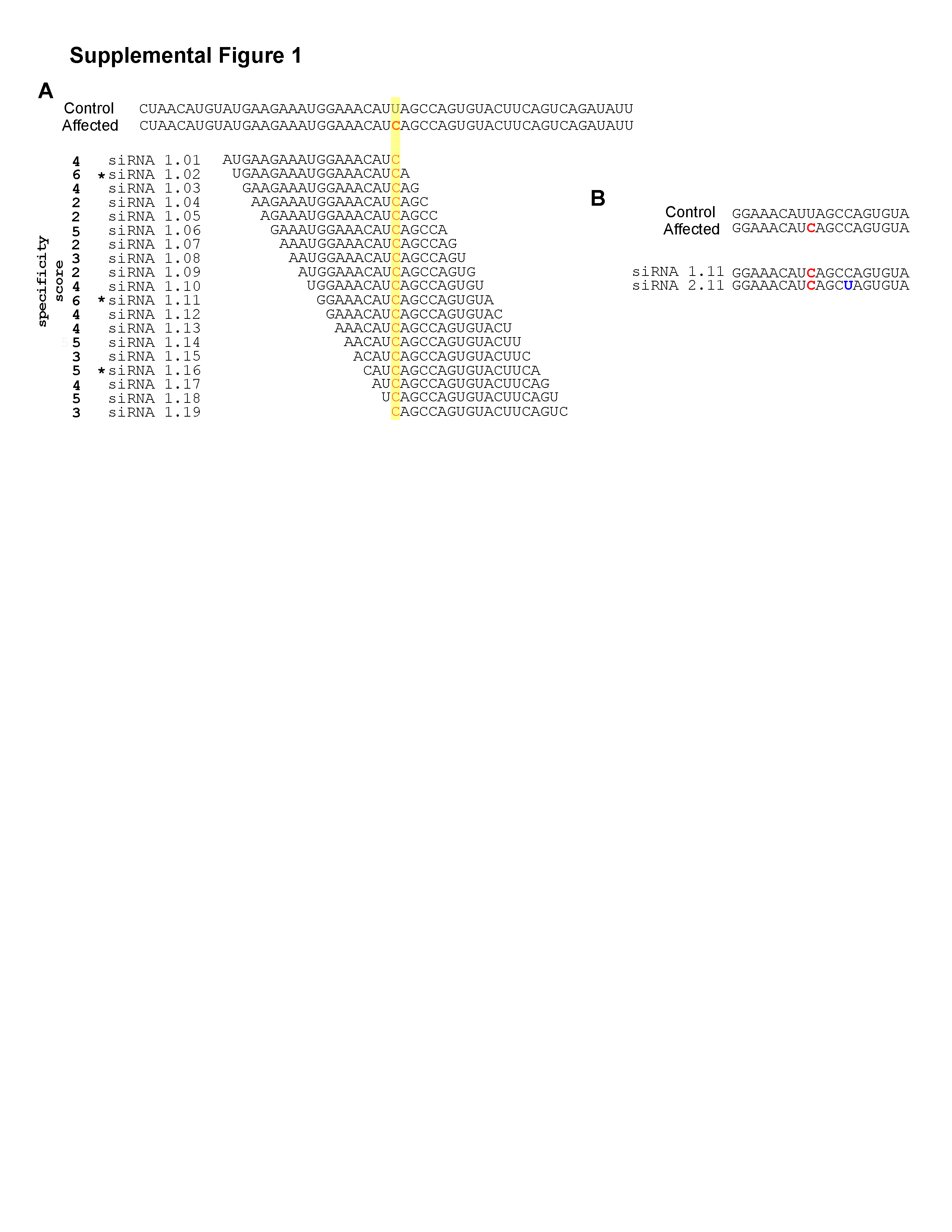

### Figure S2

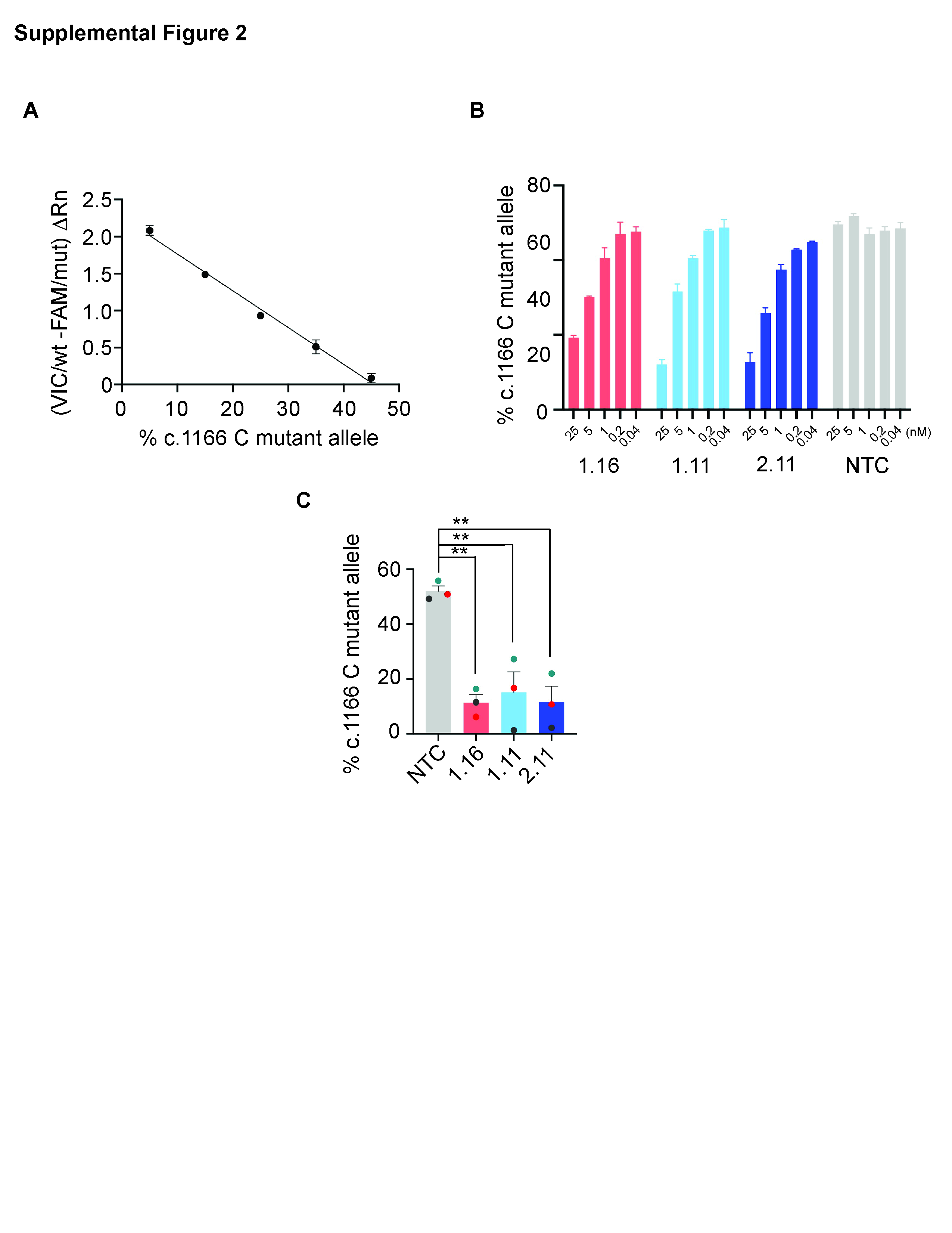

### Figure S3

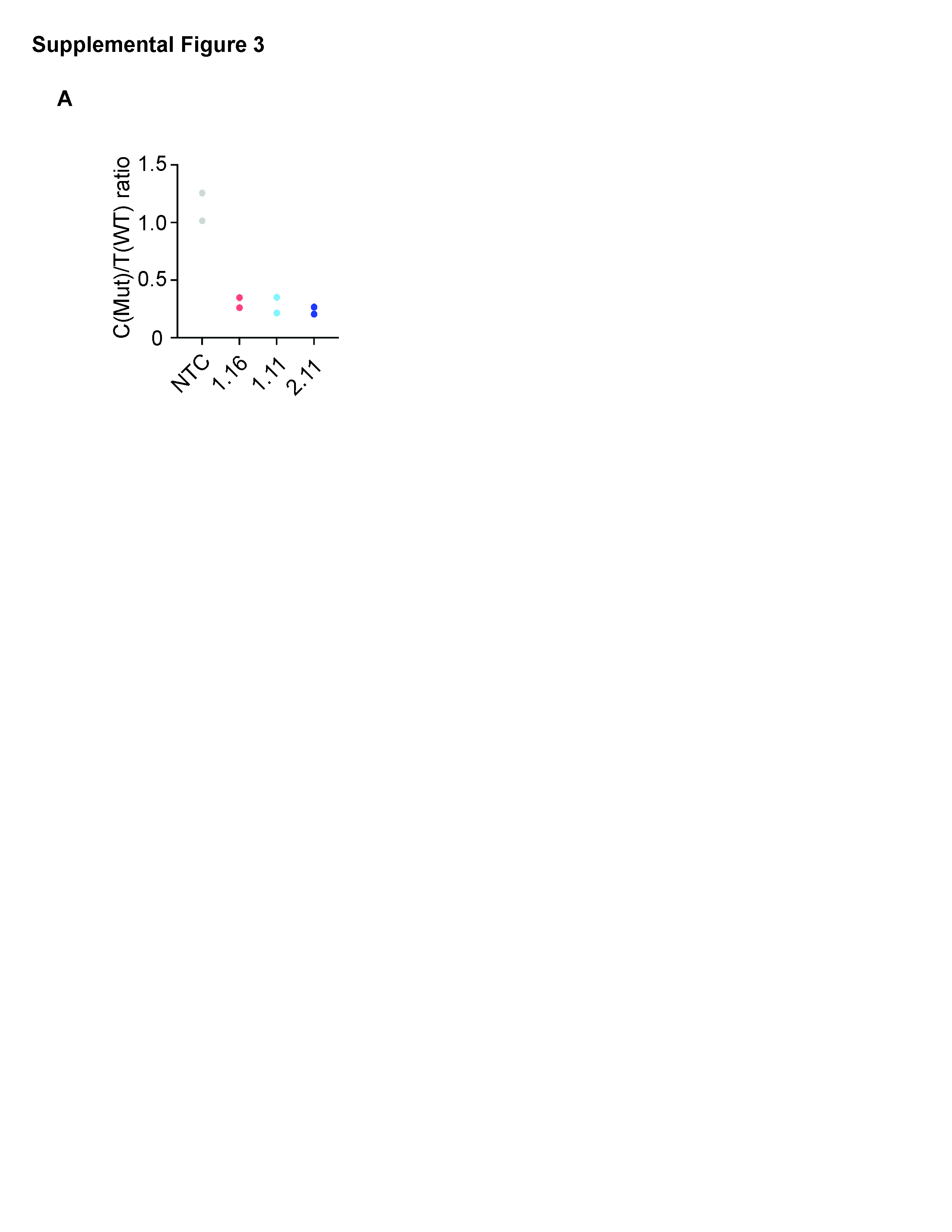
